# “Helicase” Activity Promoted Through Dynamic Interactions Between a ssDNA Translocase and a Diffusing SSB Protein

**DOI:** 10.1101/2022.09.30.510372

**Authors:** Kacey Mersch, Joshua E. Sokoloski, Binh Nguyen, Roberto Galletto, Timothy M. Lohman

**Author notes:** These authors contributed equally to this work (Kacey Mersch, Joshua E. Sokoloski).

## Abstract

Replication protein A (RPA) is a eukaryotic single stranded (ss) DNA binding (SSB) protein that is essential for all aspects of genome maintenance. RPA binds ssDNA with high affinity but can also diffuse along ssDNA. By itself, RPA is capable of transiently disrupting short regions of duplex DNA by diffusing from a ssDNA that flanks the duplex DNA. Using single molecule total internal reflection fluorescence and optical trapping combined with fluorescence approaches we show that *S. cerevisiae* Pif1 can use its ATP-dependent 5’ to 3’ translocase activity to chemo-mechanically push a single human RPA (hRPA) directionally along ssDNA at rates comparable to those of Pif1 translocation alone. We further show that using its translocation activity Pif1 can push hRPA from a ssDNA loading site into a duplex DNA causing stable disruption of at least 9 bp of duplex DNA. These results highlight the dynamic nature of hRPA enabling it to be readily reorganized even when bound tightly to ssDNA and demonstrate a new mechanism by which directional DNA unwinding can be achieved through the combined action of a ssDNA translocase that pushes an SSB protein.

## Introduction

Single stranded (ss) DNA binding (SSB) proteins are essential for all aspects of genome maintenance. They function by binding to ssDNA formed transiently during DNA replication, recombination and repair to protect the strands from chemical and enzymatic attack, to disrupt the formation of secondary structure in the unpaired ssDNA, and to recruit other proteins and enzymes to their sites of action. In eukaryotes, the primary SSB is replication protein A (RPA), a hetero-trimer with multiple oligosaccharide/oligonucleotide binding (OB) fold domains. Human (h) RPA consists of a 70 kDa Rpa1 subunit containing 4 OB folds (F, A, B, and C), a 32 kDa Rpa2 subunit with 1 OB fold (D), and a 14 kDa Rpa3 subunit with 1 OB fold (E) (1, 2). Although hRPA binds ssDNA with quite high affinity, its DNA binding is dynamic (1, 3, 4). hRPA can diffuse along ssDNA and this diffusion is central to the mechanism by which hRPA can transiently disrupt DNA secondary structure (hairpins) that it encounters along the ssDNA (3, 5), similar to the activity observed for *E. coli* SSB protein (6, 7).

During DNA replication, recombination, and repair, SSB-ssDNA complexes form transiently since their displacement or transfer from ssDNA must occur before dsDNA can be reformed. Such reorganization of SSB-ssDNA complexes can occur by mechanisms such as complex dissociation or direct transfer to another segment of ssDNA (8, 9) or through active SSB exchange (10, 11). We demonstrated that superfamily (SF) 1 ssDNA translocases are able to chemo-mechanically push *E. coli* SSB tetramers along ssDNA using their ATP-dependent directional motor activity (12). It has also been shown that ssDNA translocases can displace RPA from ssDNA (13).

Using two complementary single molecule approaches (smTIRF microscopy and dual optical-trapping combined with fluorescence confocal microscopy), we report that hRPA can be pushed along ssDNA by the chemo-mechanical action of the *Sc*Pif1 ssDNA translocase in the same manner as *E. coli* SSB tetramers (12), even though these two SSBs are structurally very different (14). We further show that under conditions in which ScPif1 does not act as a helicase, it can use its ssDNA translocase activity to push hRPA directionally into duplex DNA, resulting in a long-lived disruption beyond the limits that hRPA could achieve by simple diffusion (3). Hence, the action of a diffusing SSB protein and a directional ssDNA translocase motor can be combined to generate a novel “helicase” activity that may function to generate ssDNA flanking regions that can be used to initiate DNA repair processes or to regulate access to the 3’ end of a ss/dsDNA junction.

## Results

### Replication Protein A diffuses along ssDNA

Using a single molecule total internal reflection fluorescence (smTIRF) microscopy assay Nguyen et al. (2014) showed that hRPA can diffuse along ssDNA. A one-dimensional diffusion coefficient on ssDNA of ∼ 3000 nt^2^/sec, was estimated for human RPA (hRPA) at 25°C (500 mM NaCl) based on modelling of the smTIRF data. Here we used a smTIRF FRET assay to demonstrate hRPA diffusion along a short 60 nucleotide ssDNA as described (3). In this case, ssDNA labeled with a Cy3 fluorophore at its 3’ end (3’-dT-Cy3-(dT)_60_) was hybridized to a biotinylated 18 bp dsDNA handle and immobilized on the surface of a neutravidin-treated biotinylated-PEG coated coverslip (**Figure 1A)**. The sequence and structures of all smTIRF DNA substrates are given in **Supplementary Table 1 and Supplementary Table 2**. smTIRF measurements were performed in Buffer A at 25°C (see Materials and Methods). Stable Cy3 emission was observed while imaging DNA alone (**Figure 1A**). When Cy5-labeled human RPA (Cy5-hRPA) (see Materials and Methods) (100 pM to 1 nM) was added to the immobilized DNA, followed by washing out unbound hRPA, the formation of Cy5-hRPA/DNA complexes resulted in the appearance of fluctuating anti-correlated Cy3 and Cy5 fluorescence changes **(Figure 1B)**. The fluctuating FRET signal results from the random diffusion of hRPA along the ssDNA such that the Cy5-hRPA transiently approaches and recedes from the Cy3 at the 3’ end of the DNA (3).

**Table 1.**
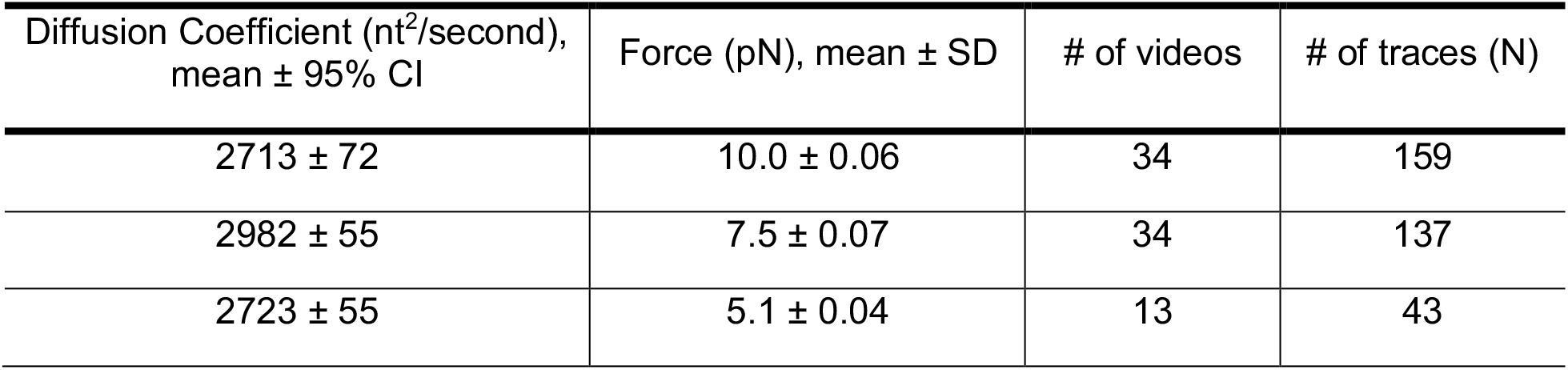
Cy5-hRPA Diffusion Coefficients.

| Diffusion Coefficient (nt<sup>2</sup>/second),<br>mean ± 95% CI | Force (pN), mean ± SD | # of videos | # of traces (N) |
| --- | --- | --- | --- |
| 2713 ± 72 | 10.0 ± 0.06 | 34 | 159 |
| 2982 ± 55 | 7.5 ± 0.07 | 34 | 137 |
| 2723 ± 55 | 5.1 ± 0.04 | 13 | 43 |

**Figure 1.**
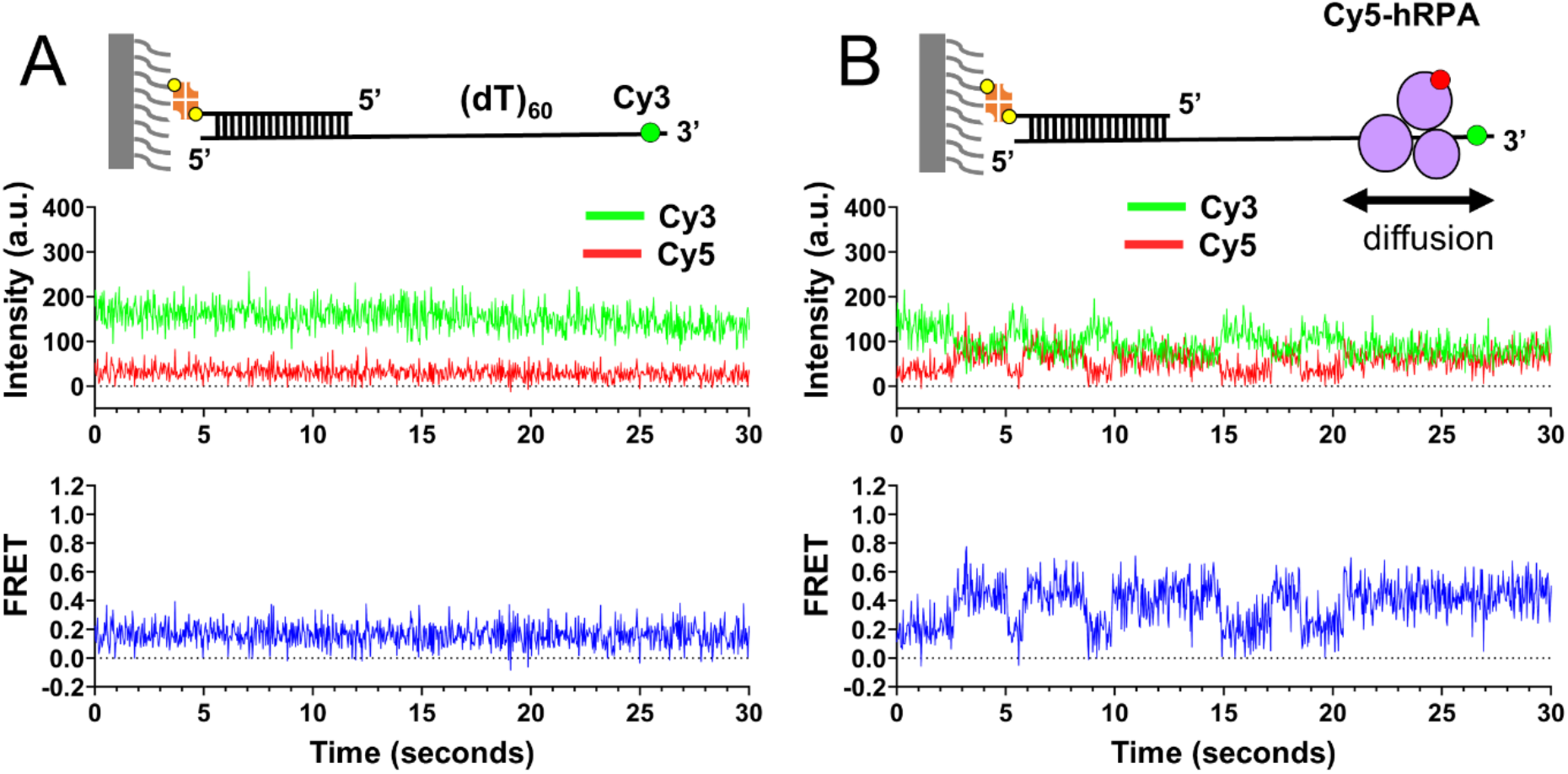
hRPA diffuses along single-stranded DNA. **A)** Representative time trajectory from a smTIRF experiment for a single molecule of 3’-Cy3-labeled (dT)_60_ ssDNA immobilized on the surface of a coverslip as shown in the cartoon. **B)** To monitor protein diffusion, a single Cy5-hRPA was bound to the surface immobilized Cy3-labeled ssDNA substrate as depicted in the cartoon. The anti-correlated Cy3 and Cy5 time trajectories and the resulting FRET fluctuations reflect the diffusion of Cy5-hRPA along ssDNA.

In order to analyze hRPA diffusion along ssDNA quantitatively and obtain an estimate of a one-dimensional diffusion coefficient, we used confocal scanning microscopy paired with an optical tweezers (LUMICKS C-Trap G2). In these experiments, the ends of a long (20,452 nt) ssDNA were held under tension between two optical traps in a Lumicks flow cell. The ssDNA tether was formed by capturing the ends between two streptavidin-coated polystyrene beads of the 20,452 bp dsDNA construct containing a 5x-biotin tag on both the 3’ and 5’ end of one strand of the DNA duplex (**Supplementary Figure 1A**). Increasing tension was applied to the dsDNA tether up to ∼82 pN and held momentarily at this force to allow the strand lacking the biotin tags to dissociate from the tethered DNA strand (**Supplementary Figure 1B**). A new force-extension profile was then obtained confirming that the tethered DNA was ssDNA.

Following generation of a ssDNA tether and binding of Cy5-hRPA **(Supplementary Figure 1A)**, the movement of Cy5-hRPA along the ssDNA was monitored by holding the ssDNA tether under tension while scanning the length of the ssDNA in 27.2 ms increments using a 638 nm confocal laser. **Figure 2A** shows a kymograph monitoring four Cy5-hRPA molecules bound to the ssDNA under a 10 pN force, imaged at 30°C in Buffer B. We obtained trajectories for 159 Cy5-hRPA molecules, and these are all displayed in **Figure 2B**, with the initial position of each trace offset to start at zero. These data show that the Cy5-hRPA trajectories adhere to simple diffusion behavior, as the traces in **Figure 2B** exhibit dynamic movements away from an initial starting point. Furthermore, the mean displacement of Cy5-hRPA molecules (orange line) follows closely along the zero position, indicating that, on average, the molecules exhibit no net displacement on ssDNA, consistent with simple Brownian diffusion.

**Figure 2.**
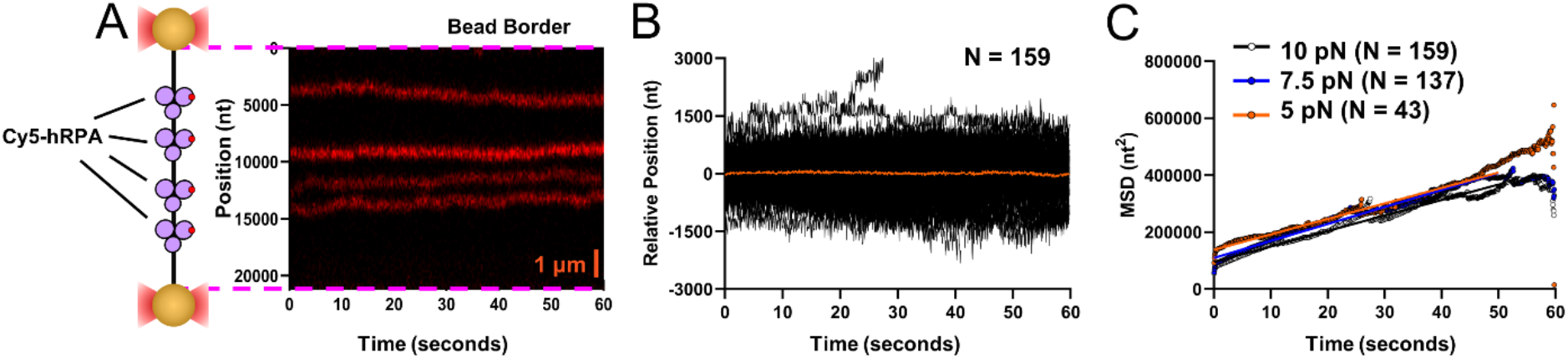
Quantification of hRPA diffusion on single-stranded DNA. **A)** Cy5-labeled hRPA was bound to ssDNA (20,452 nucleotides) that was immobilized between two streptavidin coated polystyrene beads and held at constant force in a Lumicks C-Trap. Kymographs of four Cy5-hRPA molecules obtained by scanning the DNA length repetitively with a 638 nm excitation laser show the diffusive movements of Cy5-hRPA along the ssDNA. **B)** Trajectories from 159 Cy5-RPA molecules are shown (each offset to begin at a position of zero) for ssDNA under 10 pN. The orange line shows that the mean displacement for the entire dataset is zero indicating that Cy5-hRPA moves stochastically along ssDNA. **C)** The Mean Squared Displacement (MSD) of Cy5-hRPA on ssDNA held at the indicated forces was fit with linear lines to 50 seconds. (See Table 1 for Diffusion Coefficients).

We obtained quantitative estimates of one-dimensional diffusion coefficients (D_1_) for Cy5-hRPA on ssDNA held at three forces (10, 7.5, and 5 pN). **Figure 2C** shows plots of the mean squared displacement (MSD) as a function of time at each force. The MSD for each trajectory was obtained using Eq. (1) (see Materials and Methods), and **Figure 2C** presents averaged MSD values obtained at each force. The data at each force were fit by linear regression and diffusion coefficients obtained from the slopes divided by 2, since MSD = 2Dt. The resulting mean diffusion coefficient estimates (**Table 1**) show little dependence on force, with an average mean one-dimensional diffusion coefficient, D_1_ = 2806±153 nt^2^/sec at 30°C.

### ScPif1 Translocase Can Push Replication Protein A along ssDNA

We next investigated the behavior of hRPA in the presence of an ATP-dependent ssDNA translocase. We wanted to examine the effects of a heterologous translocase and RPA pair so that there would be no specific interactions between the two proteins, hence we chose the *Saccharomyces cerevisiae* (Sc) Pif1 and the human RPA protein. We first performed smTIRF experiments to observe ScPif1 translocation and pushing of hRPA using the short (dT)_60_ ssDNA (3’-dT-Cy3-(dT)_60_) described above. Upon addition of unlabeled ScPif1 and ATP in Buffer A we observed the appearance of time trajectories showing rapid Cy3 fluorescence fluctuations (**Figure 3A**). These Cy3 fluctuations result from repetitive ATP-dependent ScPif1 translocation events on ssDNA (12, 15). A Cy3 enhancement occurs when unlabeled ScPif1 encounters the Cy3 fluorophore at the 3’ end of the (dT)_60_ due to protein-induced Cy3 fluorescence enhancement (PIFE) (16, 17). Because Pif1 preferentially binds to the ds/ssDNA junction, translocation of Pif1 to the 3’ end results in ssDNA looping (15). After reaching the 3’ end, Pif1 releases the 3’ end while remaining bound to the ss/ds DNA junction, thereby resulting in repetitive translocation cycles and repetitive Cy3 enhancements (15).

**Figure 3.**
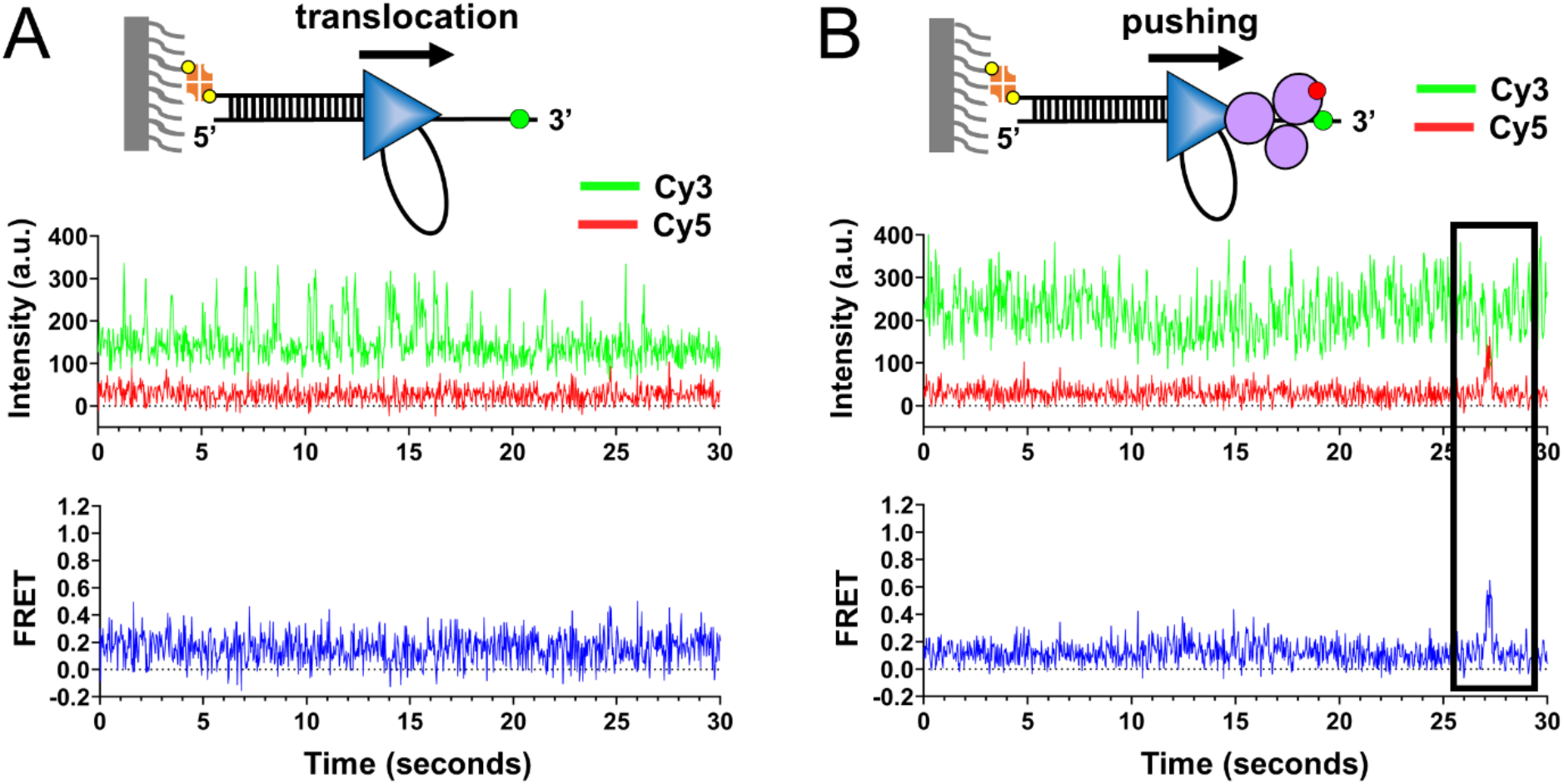
Translocating Pif1 pushes hRPA off single-stranded DNA. **A)** Representative time trajectory from a smTIRF experiment using a 3’ Cy3-labeled (dT)_60_ ssDNA substrate immobilized on the surface of a coverslip in the presence of 100 nM Pif1 (schematically depicted in the cartoon as a blue triangle) and 5 mM ATP. Spikes of enhancement in Cy3 within the Cy3 time trajectory can be observed as Pif1 comes in close contact with Cy3. **B)** Before the addition of Pif1 and ATP, 100 pM Cy5-hRPA (shown in the cartoon as magenta circles) was added to the DNA, and unbound protein was removed by washing. The anti-correlated changes in the Cy3 and Cy5 time trajectories and the associated FRET spike (boxed in black) result from Cy5-hRPA being pushed off the end of the Cy3-labeled ssDNA by Pif1.

The same experiment was then performed by first adding Cy5-hRPA followed by unlabeled ScPif1 plus ATP. In this case, we observed a decrease in the number of time trajectories displaying the FRET signals associated with Cy5-RPA diffusion, and a concomitant increase in the Cy3 fluctuations associated with repetitive ScPif1 translocation. However, in the midst of these rapid Cy3 fluorescence fluctuations, we observe occasional isolated asymmetric Cy5 fluorescence increases and concomitant Cy3 fluorescence decreases resulting in a FRET spike **(Figure 3B and Supplemental Figure 2B & 2C)**. These events are similar to those observed in a previous study of *E. coli* SSB protein and ScPif1 (12). These asymmetric FRET spikes indicate that Cy5-hRPA is being chemo-mechanically pushed by the 5’ to 3’ translocase activity of ScPif1 towards the 3’ end and eventually displaced from it. No FRET spikes are observed in the absence of ATP. Furthermore, the time it takes for the signal to rise from the zero FRET baseline to the peak FRET signal decreases with increasing ATP concentration indicating a coupling of these events to the ScPif1 ATPase activity **(Supplemental Figure 1D and 1E)** (12, 18).

To quantitatively examine ScPif1 pushing of hRPA along ssDNA, we again utilized a Lumicks C-Trap equipped with a scanning confocal laser to observe the movement of Cy5-hRPA along the 20,452 nucleotide long ssDNA in the presence of ScPif1 and ATP **(Figure 4A)**. Experiments were performed by moving the ssDNA tether held under force (10 pN) with bound Cy5-hRPA in Buffer B into a channel containing 100 nM ScPif1 and 5 mM ATP in Buffer B at 30°C **(Supplementary Figure 1A)**. Once moved into the channel containing ScPif1 and ATP, the Cy5-hRPA molecules showed uni-directional movement towards one end of the ssDNA (red kymograph in **Figure 4A)**, presumably the 3’ end since ScPif1 is a 5’-to-3’ ssDNA translocase (18). Each Cy5-hRPA was tracked until it reached the polystyrene bead or until the signal disappeared indicating either photobleaching or dissociation of hRPA. Each kymograph was fit by linear regression to obtain a pushing rate of 310 ± 76 nt/s (n = 89) at 5 mM ATP and 184 ± 77 nt/s (n = 78) at 0.5 mM ATP **(Figure 4C)**. The slower rate at 0.5 mM ATP indicates the directional movement of Cy5-hRPA is dependent on the ATPase activity of the ScPif1 translocase.

**Figure 4.**
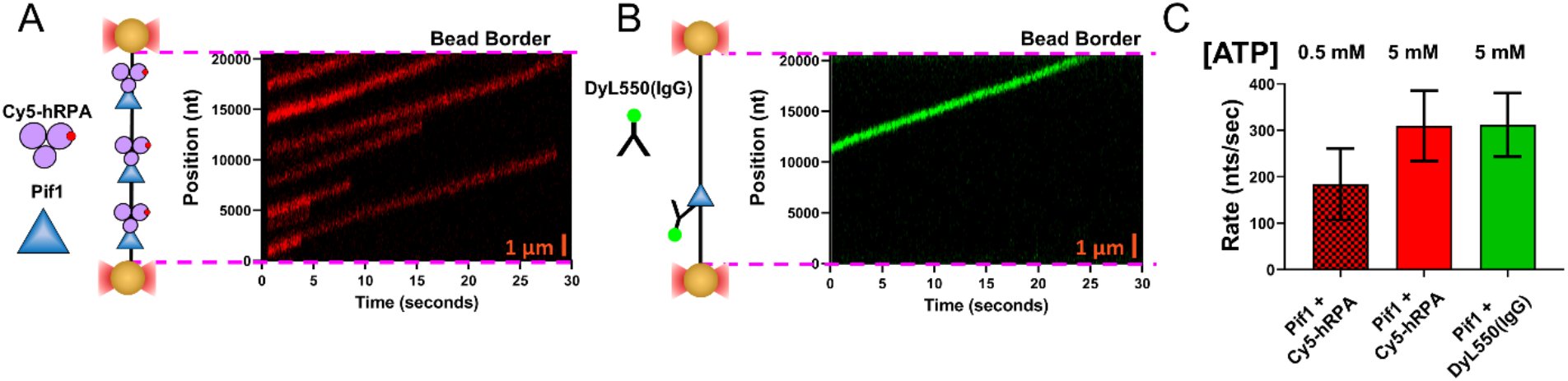
Translocating Pif1 pushes hRPA along single-stranded DNA. **A)** Cy5-hRPA was bound to a ssDNA tether as described in Figure 2A, followed by incubation with 100 nM Pif1 and 5 mM ATP. Kymographs of several Cy5-hRPA molecules (scanning the length of the DNA repetitively with a 638 nm laser) show Pif1-dependent directional movement of Cy5-hRPA along the ssDNA **B)** Translocation of Pif1 alone was monitored by binding DyL550(IgG)-labeled Pif1 to DNA in the same configuration as A, followed by incubation with 5 mM ATP. Kymograph of a representative DyL550(IgG)-Pif1 moving directionally along the ssDNA tether. **C)** Average rates of the Pif1 pushing Cy5-hRPA in the presence of 5 mM ATP (red, N = 89) or 0.5 mM ATP (red and black checker, N = 78). The average translocation rate of DyL550(IgG)-labeled Pif1 alone was measured in the presence of 5 mM ATP (green, N = 26).

We also determined the translocation rate of ScPif1 by itself in the absence of pushing Cy5-hRPA by utilizing a ScPif1 variant containing an N-terminal 6x-His tag. The ScPif1 variant was labeled by binding it to an anti-6x-His Tag Monoclonal Antibody conjugated to a DyLight^™^ 550 Fluorophore (DyL550(IgG)). The DyL550(IgG)-ScPif1 was then bound to the ssDNA tether in Buffer B before being moved to a channel containing Buffer B plus 5 mM ATP **(Supplementary Figure 1A)**. A kymograph (green) showing the directional movement of the DyL550(IgG)-ScPif1 along the ssDNA is shown in **Figure 4B**. The DyL550(IgG)-ScPif1 kymographs were analyzed by linear regression yielding a translocation rate of 312 ± 68 nt/s (n = 26) at 5 mM ATP (**Figure 4C**). Interestingly, at this saturating ATP concentration the rate of DyL550(IgG)-ScPif1 translocating unhindered is identical to the rate for ScPif1 translocation while pushing Cy5-RPA.

### ScPif1 can push hRPA into duplex DNA causing stable base pair disruption

Nguyen et al. (2014) (3) showed that hRPA is able to transiently disrupt a few bp of a duplex DNA hairpin by diffusing into the hairpin from an adjacent ssDNA loading site. However, the results indicated that the ability for hRPA to disrupt a hairpin is limited to less than ∼ 9 bp of a 18 bp hairpin (3). Since we have shown here that ScPif1 can push hRPA along ssDNA, we investigated whether ScPif1 can apply a continuous force to hRPA that can cause a stable disruption of a DNA hairpin beyond the limit observed with hRPA alone.

To examine hairpin disruption, we designed a DNA containing a (dT)_45_ -ssDNA region with an 18 bp hairpin at the 3’ end that contains a Cy3 label located in the middle of the hairpin duplex, 9 bp from both the loop and the ss/dsDNA junction **(Figure 5A)**. All smTIRF experiments were performed in Buffer A with a 532 nm laser to excite Cy3 fluorescence. Upon addition of 100 pM Cy5-hRPA to this surface bound DNA and subsequently washing out unbound Cy5-hRPA, no changes in Cy3 or Cy5 signals are observed **(Figure 5A)** indicating that Cy5-hRPA alone is unable to disrupt enough of the duplex DNA to reach the Cy3 fluorophore and illicit a FRET signal. However, upon addition of ScPif1 (100 pM) and 50 μM ATP to the Cy5-hRPA/DNA complex, we observe time trajectories with long-lived high FRET signals **(Figure 5B)** indicating that ScPif1 can push the Cy5-hRPA causing it to disrupt the duplex and reach the Cy3 fluorophore located 9 bp internally within the duplex DNA. The high FRET signals observed are stable for long periods of time (durations range from 5 sec to greater than 35 sec). These data indicate that ScPif1 can push Cy5-hRPA into the duplex DNA to reach the Cy3 and that Cy5-hRPA remains near the Cy3 before either dissociating or returning to the ssDNA region.

**Figure 5.**
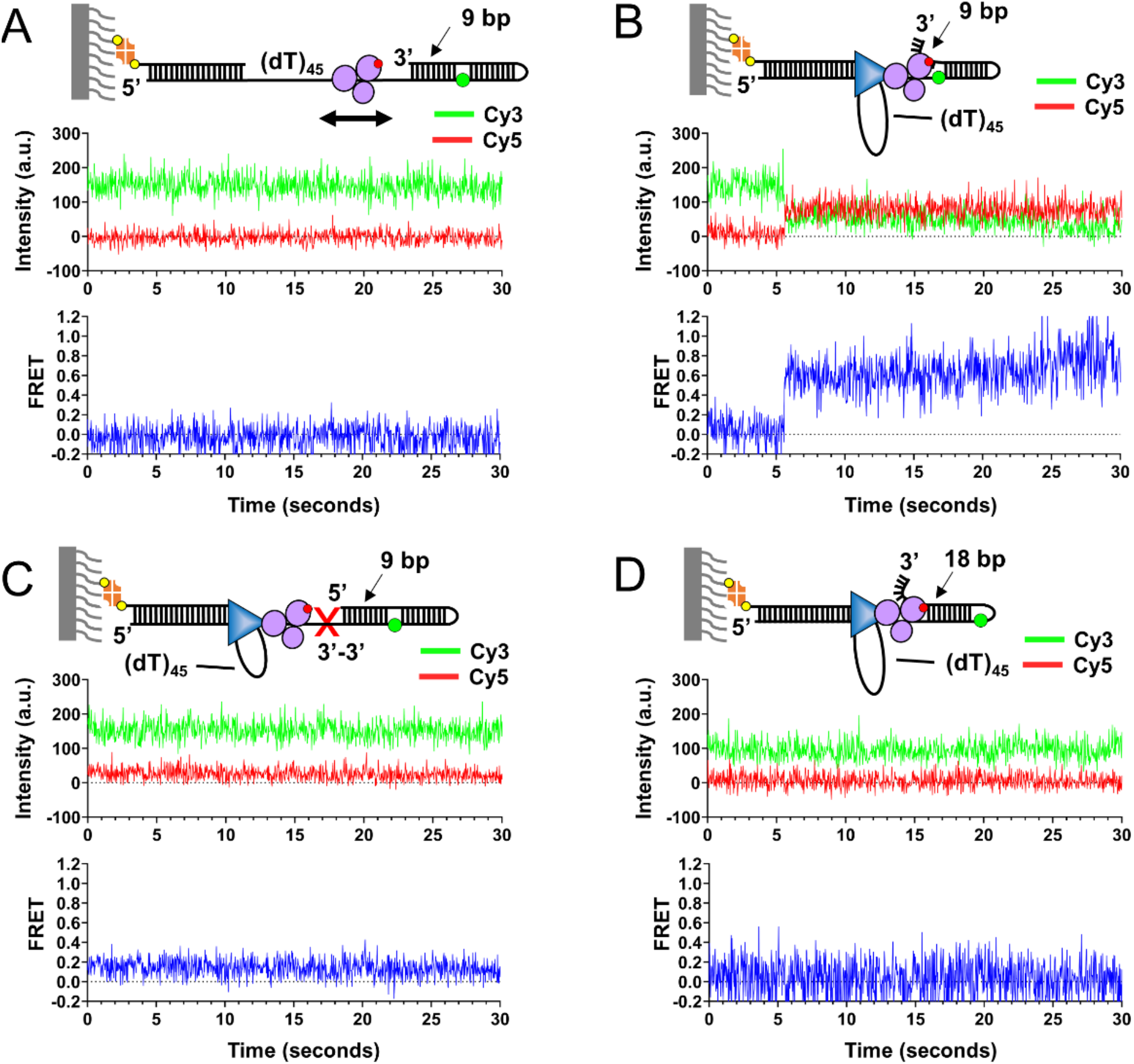
Pif1 pushes Cy5-hRPA to stably disrupt a 9 bp duplex. **A)** Cy5-hRPA was bound to a surface immobilized DNA with a (dT)_45_ ssDNA region followed at the 3’ end by an 18 bp hairpin. As schematically depicted in the cartoon, a Cy3 is placed in the backbone at 9 bp from the ss/dsDNA junction. No FRET events were observed after incubation with 100 pM of Cy5-hRPA, followed by removal of unbound Cy5-hRPA (n = 0, N = 770, 0%). **B)** After 100 pM Pif1 and 50 μM ATP are added to Cy5-hRPA/DNA complexes, high FRET signals appear, indicating that Cy5-hRPA is closer to the Cy3 fluorophore located in the center of the hairpin. FRET events were observed in ∼27% of the single molecule trajectories (n = 196, N = 714). **C)** DNA substrate similar to A, but with a reverse polarity linkage between the (dT)_45_ linker and 18 bp hairpin. No FRET signals occur in either the presence of Cy5-hRPA alone (n = 0, N = 582) or Cy5-hRPA in the presence of 100 pM Pif1 and 5 mM ATP (n = 0, N = 984). **D)** When the Cy3 was placed in the backbone at 18 bp from the ss/dsDNA junction (instead of 9 bp as in A-C) FRET events were not observed in either the presence of Cy5-hRPA (n = 0, N = 189) or the presence of 100 pM Pif1 and 50 μM ATP (n = 0, N = 285).

As a control, we tested whether ScPif1 could unwind the 5’-ssDNA tailed hairpin DNA in the absence of Cy5-hRPA at the Pif1 and ATP concentrations at which Cy5-hRPA is pushed into the duplex DNA **(Supplemental Figure 3)**. In the absence of ScPif1, a stable Cy3 fluorescence signal is observed for DNA alone **(Supplemental Figures 3A and 3C)**. Upon addition of high concentrations of ScPif1 (≥ 1 nM) and high [ATP] (5 mM), significant Cy3 fluorescence fluctuations (Cy3-PIFE signals) are observed indicating that ScPif1 can unwind the DNA and reach the Cy3 fluorophore **(Supplemental Figures 3B and 3C)**. However, no such Cy3 PIFE signals were observed at a ScPif1 concentration of 100 pM and 50 μM ATP **(Supplementary Figure 3C)**, indicating that ScPif1 alone cannot unwind the DNA to reach the Cy3 at the concentrations used in the RPA pushing experiments. This is consistent with the observation that a monomer of ScPif1 cannot unwind a dsDNA with only a single 5’ flanking ssDNA (19, 20).Thus, the high sustained FRET events observed in **Figure 5B** with Cy5-hRPA and 100 pM ScPif1 and 50 μM ATP are due to Cy5-RPA being pushed into the duplex by the translocase activity of ScPif1, since at these conditions no DNA unwinding (helicase) activity is detected with ScPif1 alone.

Furthermore, the absence of a sharp Cy3 fluorescence increase before the steady rise in FRET indicates that ScPif1 is translocating behind and pushing Cy5-hRPA rather than ScPif1 first unwinding the DNA followed by hRPA binding to the newly generated ssDNA behind the ScPif1 helicase. In support of this conclusion, when the ssDNA contains a 3’-3’ reverse polarity phosphodiester linkage at the ss/dsDNA junction, no sustained high FRET signals are observed in the presence of bound Cy5-hRPA or with the addition of 100 pM ScPif1 and 5 mM ATP **(Figure 5C)**. This indicates that neither ScPif1 nor Cy5-hRPA can proceed past the reverse polarity linkage. Hence hRPA access to the hairpin is from the adjacent ssDNA. When these experiments were repeated with a DNA substrate in which the Cy3 was placed 18 bp from the ss/ds DNA junction, no FRET signals were observed in the presence of bound Cy5-hRPA and 100 pM ScPif1 and 50 μM ATP **(Figure 5D)** or in the presence of bound Cy5-hRPA alone. Thus, the ability of ScPif1 to push Cy5-hRPA stably into the DNA duplex is limited to between 9 -18 bp.

## Discussion

The complementary single molecule approaches used here show that hRPA protein is highly dynamic; it can diffuse along ssDNA and be pushed directionally by the ATPase activity of a ssDNA translocase, ScPif1. We also report direct estimates of the hRPA diffusion coefficient on ssDNA, the rate of ScPif1 translocation and the rate of ScPif1 pushing of hRPA along ssDNA using the Lumicks C-Trap. Finally, we demonstrate a new mechanism for how hRPA, when combined with ScPif1, can stably disrupt at least 9 bp of a duplex DNA as a result of hRPA being physically pushed into the duplex DNA by the SF1 translocase. These results highlight the dynamic nature of hRPA, and presumably RPAs from other organisms. Thus RPA can be readily reorganized, even when bound tightly to ssDNA.

Previous studies have shown that *E. coli* SSB proteins (6) and eukaryotic RPA proteins (3, 5) are able to diffuse along short ssDNA. Here, we observed the diffusive movements of hRPA directly on long ssDNA by tracking a fluorescently labeled hRPA along a ssDNA under tension in an optical trap. This approach enabled a direct estimate of the one-dimensional diffusion coefficient of hRPA on ssDNA, with D_1_ ∼2700-2900 nt^2^/s at 30°C (pH 8.1, 100 mM NaCl, 5 mM MgCl_2_). Diffusion measurements were performed with the ssDNA held at forces from 5-10 pN, tensions that are sufficient to destabilize DNA secondary structures that may be present at lower forces (21, 22). The measured diffusion coefficients, which show no dependence on force in this range are in reasonable agreement with the previous estimate for hRPA of D_1_ = 2800 ± 200 nt^2^/s at 25°C (pH 8.1, 500 mM NaCl), with an extrapolated value of D_1_ = 3600 ± 300 nt^2^/sec at 30°C (3) using a smTIRF approach on short oligo(dT) DNA. The differences in these estimates could be due to the different solution conditions used and possibly because the estimates of Nguyen et al. (3) were indirect and required modelling of the smTIRF data. At the lower salt concentration used in the current study (0.1 M NaCl, 5 mM MgCl_2_), hRPA should bind to ssDNA with higher affinity than at the 0.5 M NaCl used previously (3), which might affect its ability to diffuse, however this is not the case. Other experimental variables, such as the base composition of the DNA or the application of force may also affect diffusion. Nonetheless, these combined studies indicate that hRPA is capable of diffusion along ssDNA over a range of solution conditions.

We further showed that ScPif1 is able to use its ATP-dependent uni-directional translocation activity to push hRPA along ssDNA held at 10 pN force at a rate of 310 ± 76 nt/s (5 mM ATP). We also find that ScPif1 on its own translocates on the ssDNA tether at the same rate as when it is pushing hRPA. Two limiting models have been proposed for how a translocating enzyme might push a diffusing protein directionally along ssDNA, a “direct pushing” model and a “moving barrier rectification” model (12). In the “direct pushing” model, a collision between ScPif1 and hRPA is perfectly inelastic, resulting in ScPif1 and hRPA moving as a single unit along ssDNA with the same velocity as ScPif1 alone (no load or friction from RPA). In the “moving barrier rectification” model, a Pif1 collision with RPA blocks the 5’ to 3’ translocation of Pif1 and also prevents RPA from taking a diffusive step in the 5’ direction; as a result, ScPif1 can only translocate towards the 3’ end of the ssDNA when RPA takes a diffusive step in the 3’ direction. Of these two, the direct pushing model provides a better description of ScPif1 pushing of hRPA since the load from pushing hRPA has no effect on the Pif1 translocation rate while the rectification model predicts that the pushing rate should be much slower than the rate of Pif1 translocation by itself (12).

Finally, our studies reveal a novel mechanism for the stable disruption of a short region of duplex DNA that involves rectification by the Pif1 translocase of hRPA’s ability to transiently disrupt a DNA duplex. Under conditions where neither hRPA nor ScPif1 alone can disrupt the duplex region of an 18 bp hairpin, their combined actions instead stably open at least 9 bp of duplex to produce a 3’ ssDNA flanking region. Biologically, this type of activity may help with the resolution of DNA secondary structures that impair replication, recombination, and repair at sites of genomic maintenance. In this regard, we note that one function of Pif1 helicases is to remove obstacles that would otherwise impede efficient progression of DNA replication. For example, Pif1 stimulates DNA replication by unwinding stable G-quadruplexes (7, 23-25), removing proteins tightly bound to DNA (26, 27) or a Cas9-dependent R-loop (28). In these cases the helicase activity of Pif1 appears to be needed to unwind the DNA in front of a DNA polymerase. However, here we show that the ssDNA translocase activity of Pif1 can also be used to destabilize a duplex DNA, by forcing an SSB protein into the duplex. Previous studies show that a single hRPA hetero-trimer cannot transiently destabilize 9 bp of an 18 bp DNA hairpin (3), whereas here we show that ScPif1 can push a single hRPA into a hairpin, disrupting at least 9 bp for times that exceed several seconds. The question remains, as to how the transient destabilization of a short duplex by RPA (3) or as shown here, the stable Pif1-dependent disruption of a longer duplex by RPA could be functional. Transient or stable opening of a short stretch of duplex DNA by these means may facilitate DNA repair processes that require the generation of a flanking 3’ ssDNA that could be used as a loading site for other proteins, such as RecA or RAD51 recombinases (29-33). Alternatively, since the 3’ end of a ss/ds junction of the duplex is a loading site for a DNA polymerase, its transient disruption by an SSB or stable opening by an SSB being pushed by a translocase could provide a means to regulate its use as a site for DNA synthesis.

We purposely used a heterologous translocase and RPA pair from different organisms so that we could examine how RPA responds to the force exerted by a directionally translocating enzyme in the absence of any direct protein-protein interactions. The results in this study expand those obtained in a previous study of translocases pushing a tetrameric bacterial SSB protein (12). That study showed monomers of the superfamily 1 (SF1) ssDNA translocases, *E. coli* UvrD, *E. coli* Rep, and ScPif1, can directionally push *E. coli* SSB tetramers along ssDNA in reactions that are coupled to ATP hydrolysis (12). Rep and UvrD pushed SSB in the 3’ to 5’ direction, whereas Pif1 pushed SSB in the 5’ to 3’ direction, based on their known ssDNA translocase directionalities. That study also concluded that a “direct pushing” model provided a good description of the pushing of *E. coli* SSB protein along ssDNA by those translocases (12). There are numerous examples of SSB proteins from both prokaryotes and eukaryotes that can form specific interactions with a cognate ssDNA translocase (30, 34, 35). It will be of interest to examine how those interactions might influence the translocase rates and activity and also the ability of the SSB to invade a DNA duplex.

The results presented here also bear generally on the mechanisms of helicases. It is now recognized that DNA unwinding (helicase) activity generally requires more than just ssDNA translocase activity (36). For example, the SF1 enzymes, Rep, UvrD, and PcrA monomers are rapid and processive ssDNA translocases, but have no detectable helicase activity by themselves (37-43). These enzymes require activation in order to function as helicases and unwind DNA. This can occur by dimerization (44, 45), removal of an auto-inhibitory sub-domain (37) or through an interaction with an accessory protein, PriC for Rep (46), MutL for UvrD (47, 48), RepD for PcrA (49). Here we demonstrate a new mechanism for how the combined action of a directional ssDNA translocase and an SSB protein can acquire DNA unwinding activity. In the absence of the translocase, a single RPA hetero-trimer can transiently destabilize a short region of a DNA duplex (3). Hence, the RPA is the factor that actively destabilizes the duplex, however, this is transient. The presence of the chemo-mechanical translocase is needed to provide a directional force preventing (rectifying) the RPA from exiting the DNA duplex. This demonstrates the basic properties of a “helicase”. One component (RPA) must destabilize the duplex DNA, while another component (ATP-dependent translocase) provides the force to prevent reformation of the duplex DNA.

Our study examined the consequences of a ssDNA translocase pushing and eventually displacing a single RPA hetero-trimeric protein. It will also be of interest to examine whether a ssDNA translocase is able to push and displace multiple RPA proteins along ssDNA. Recent studies have shown that the human DNA helicase B (HelB) can use its 5’ to 3’ ssDNA translocase activity to displace multiple hRPA proteins from ssDNA (13). In that case, HelB appears to interact specifically with hRPA and this interaction stimulates the translocase activity of HelB. No stimulation was observed with the heterologous yeast RPA. In fact multiple yeast RPA proteins appeared to prevent HelB translocation along ssDNA. Although no evidence was presented concerning whether HelB can push the hRPA in those studies, it seems likely that RPA pushing would precede RPA displacement.

## Materials and Methods

### Buffers

Buffer A is 30 mM TRIS (pH 8.1), 100 mM NaCl, 5 mM MgCl_2_, 0.1 mM Na_2_EDTA, 1 mM dithiothreitol, 0.5% (w/v) dextrose, 0.1 mg/mL bovine serum albumin, 4 mM 6-Hydroxy-2,5,7,8-tetramethylchroman-2-carboxylic acid (Trolox). Buffer B is 30 mM TRIS (pH 8.1), 100 mM NaCl, 5 mM MgCl_2_, 0.1 mM Na_2_EDTA, 1 mM dithiothreitol, 0.5% (w/v) dextrose, 4 mM 6-Hydroxy-2,5,7,8-tetramethylchroman-2-carboxylic acid (Trolox). PBS buffer is 137 mM NaCl, 2.7 mM KCl, 1.19 mM Phosphates, 500 μM EDTA and 5 mM NaN_3_. Buffer X is 10 mM TRIS (pH 8.1), 50 mM NaCl, and 0.1 mM Na_2_EDTA.

### Oligodeoxynucleotide synthesi

Oligodeoxynucleotides were synthesized on a MerMade 4 synthesizer (Bioautomation, Plano, TX) using phosphoramidites (Glen Research, Sterling, VA). DNA Purification and concentration was performed as previously described (3). DNA duplexes were annealed in Buffer X as previously described (3). The sequences and structures of the oligodeoxynucleotides used in this study are shown in **Supplemental Tables 1 and 2**.

### Protein purification and labeling

Human RPA (hRPA), *S. cerevisiae* Pif1, and *S. cerevisiae* Pif1 N-His were purified and concentrations were determined by absorbance as previously described (3, 19, 50). hRPA was labeled stochastically as previously described by (Nguyen et al. 2014) (3) with Cy5 mono NHS ester (PA15101, Amersham, Piscataway, NJ) under conditions where the N-terminal amines are preferentially labeled. Labeling percentage as measured by UV-VIS absorbance was 92% (dye to hRPA ratio), indicating that while the hRPA was labeled stochastically among the three N-termini, there was only one Cy5 per hRPA. Labeling occurred on 46% of Rpa1, 34.5% of Rpa2, and 11.5% of Rpa3 according to SDS-PAGE. Labeling percentage of the Cy5-hRPA used in the optical tweezers experiments was 148% (dye to hRPA ratio). The ScPif1-N-His protein was labeled by binding to a 6x-His Tag Monoclonal Antibody (500 nM) conjugated to a DyLight^™^ 550 Fluorophore (Cat. # MA1-21315-D550, ThermoFisher Scientific).

### Single Molecule Total Internal Reflection Fluorescence Microscopy (smTIRFM)

smTIRFM experiments were conducted on an objective-based total internal reflection fluorescence microscope custom built from an IX71 inverted microscope (Olympus) with an 60x 1.45 numerical aperture PlanApo objective (Olympus) as described (3, 12). Briefly, biotinylated DNA substrates containing Cy3 were immobilized onto a polyethylene glycol (PEG, MW 5000) coated coverslip via Neutravidin linkage to biotinylated PEG. The immobilized Cy3 substrates were excited by illumination from a 532 nm diode-pumped solid state laser (CrystaLaser, Reno, NV) coupled to the microscope using KineFlex fiber optics (QiOptics). All experiments were done at 25°C using a temperature controlled stage (BC-110 Bionomic controller, 20/20 Technology Inc) and an objective heater (Bioptechs Inc). Fluorescence emission was detected with an Andor iXon EMCCD camera (Model DU897E) and SINGLE, a custom program provided by Taekjip Ha (Johns Hopkins University), was used for movie collection. Movies were processed using custom scripts in IDL (Exelis VIS) to extract individual intensity vs time trajectories and then analyzed with MatLab (Mathworks). smTIRFM experiments were performed in Buffer A and ATP concentrations as indicated for each experiment. An oxygen scavenging system (glucose oxidase (1 mg/mL final concentration) and catalase (0.4 mg/mL final concentration)) was added to all samples immediately before loading onto the TIRF slide.

### Combined Optical Tweezer and Confocal Scanning

Diffusion of Cy5-hRPA, ATP-dependent translocation of ScPif1 and pushing of hRPA by ScPif1 was performed with a LUMICKS C-Trap controlled with Bluelake™ (v2.0) software. The combined optical tweezer and confocal scanning microscope is outfitted with a μ-Flux™ Microfluidics System (LUMICKS) with a flow cell containing 5 channels (C1, LUMICKS). The LUMICKS C-Trap flow cell was passivated prior to performing experiments by flowing PBS (0.5 mL at 1.6 bar) through the syringes, lines, and flow cell. Followed by flowing 0.5% (w/v) Pluronic acid (0.5 mL at 1.6 bar, SKU00003, LUMICKS) and then 0.1% (w/v) BSA (0.5 mL at 1.6 bar, SKU00003, LUMICKS). The passivation procedure is ended by flowing PBS (0.5 mL at 1.6 bar) through the same system. Please refer to **Supplemental Figure 1** to aid in the description below. Experiments begin by procuring streptavidin coated polystyrene beads (4.89 um, SKU00003, LUMICKS), in channel 1, within each of the two optical trapping lasers (1064 nm). The beads within the traps are moved to channel 2 to bind a single 20,452 bp biotinylated DNA (SKU00014, LUMICKS) to both beads. The duplex DNA tether is moved to channel 3 where a force is applied to stretch the DNA until the unattached DNA strand dissociates yielding a 20,452 nucleotide long ssDNA tether assembly. ssDNA formation was performed in a 1/10^th^ dilution of PBS (SKU00003, LUMICKS) and was confirmed to be ssDNA through visual confirmation on the screen that no hysteresis occurred during DNA extensions. Force applied to the ssDNA tether was measured by Trap 2, while Trap 1 was moved to manipulate force on the DNA. Protein-DNA complexes were formed in channel 4, and then imaged in channel 5. Refer to **Supplemental Figure 1** for more details.

Experiments quantifying the diffusional movements of Cy5-hRPA on ssDNA were performed using a feedback loop that maintains a constant force on the ssDNA. The temperature was regulated at 30°C by temperature controllers on both the objective and condenser of the microscope. Diffusion measurements were performed using Buffer B. Glucose oxidase (0.5 mg/mL final concentration) and catalase (0.2 mg/mL final concentration) were added to the solution just before placing the mixed solutions into the microfluidics system. A freshly prepared solution was placed into the microfluidics system every two hours of image acquisition. Kymographs were acquired using a 638 nm laser at 4% power, 27.2 ms scanning line time, and a pixel size of 100 nm. Tracking of Cy5-hRPA molecules, mean squared displacement (MSD) of hRPA, and single molecule diffusion coefficients were calculated in Pylake (v0.12.1, LUMICKS) using python scripts (v3.10.5) executed within Jupyter Notebooks. Individual molecule kymograph images were obtained by implementing the greedy tracking algorithm, which finds pixels with intensities above a threshold and subsequently refines an area of interest to determine its subpixel position before linking positions on the same trajectory together to follow the position of the molecule over time. To enable more robust tracking, the time along the kymograph image was binned by two.

The mean squared displacement of Cy5-hRPA for each trace was calculated using Equation (1).

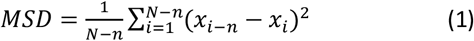

Where *n* = lag number, N = number of points within the tracked line, and *x*, = trace position at time frame i. The diffusion coefficients from the mean MSD were calculated by fitting the mean data from zero to 50 seconds using linear regression. The slopes of those best-fit lines were divided by two to obtain the mean diffusion coefficient (MSD = 2Dt).

Tracking and imaging of molecules during the Pif1 translocation experiments were performed as described for the diffusion experiments. Solution conditions were identical to those of the diffusion experiments, except that ATP was added in the translocation experiments. The tracked lines from the kymographs exhibiting pushing or translocation were placed into GraphPad Prism and fit by linear regression to measure the translocation rate.

## Acknowledgements

We thank Thang Ho for synthesis and purification of the oligodeoxynucleotides. This work was supported by National Institutes of Health Grants 5R35GM136632 (to T.M.L.) and 1R35GM139508 (to R.G.) and American Cancer Society Grant PF-15-040-01-DMC (to J.E.S). The Lumicks C-Trap G2 was purchased with support from NIH Instrumentation grant S10OD030315.

## Figures and Tables

**Figure 6.**
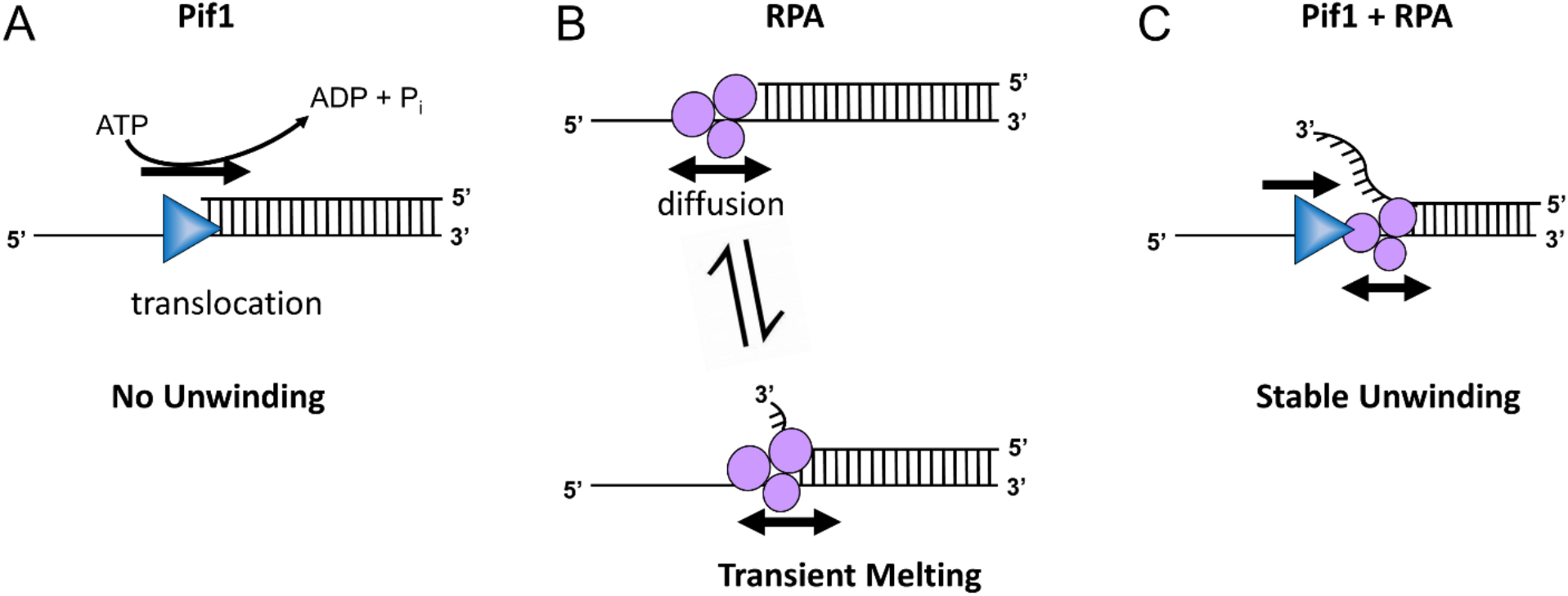
RPA diffusion rectified by Pif1 translocation leads to stable DNA unwinding. **A)** A Pif1 monomer can translocate along ssDNA, but it does not exhibit helicase activity on a single tailed dsDNA. **B)** RPA diffuses along ssDNA and can transiently melt a short region of duplex DNA. **C)** When combined, the Pif1 translocase can rectify the transient DNA melting by RPA to stably unwind at least 9 bp of duplex DNA by applying a directional force to RPA at the junction.

**Table S1.**
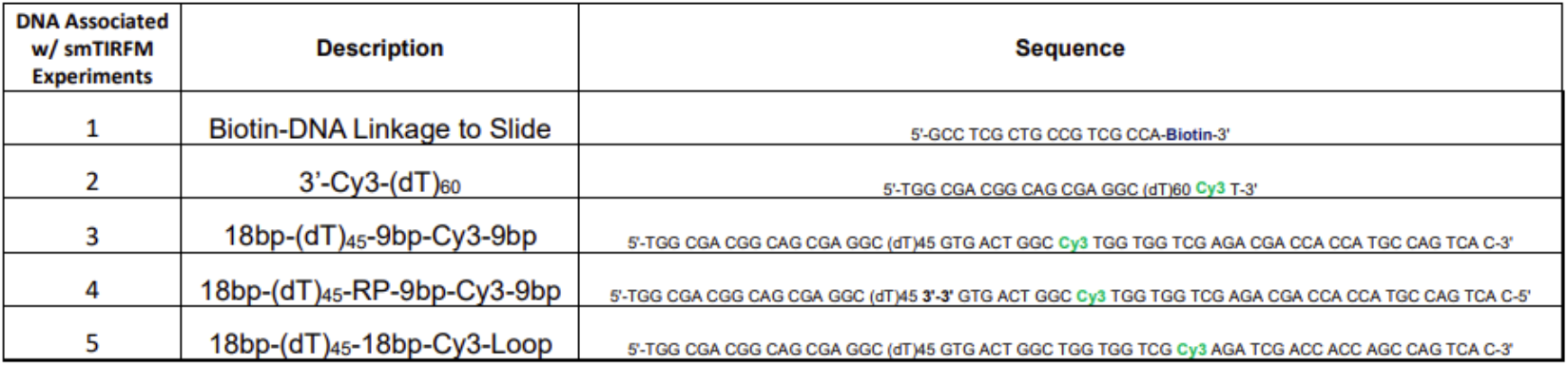
Single Molecule TIRF DNA Sequence Composition

**Table S2.**
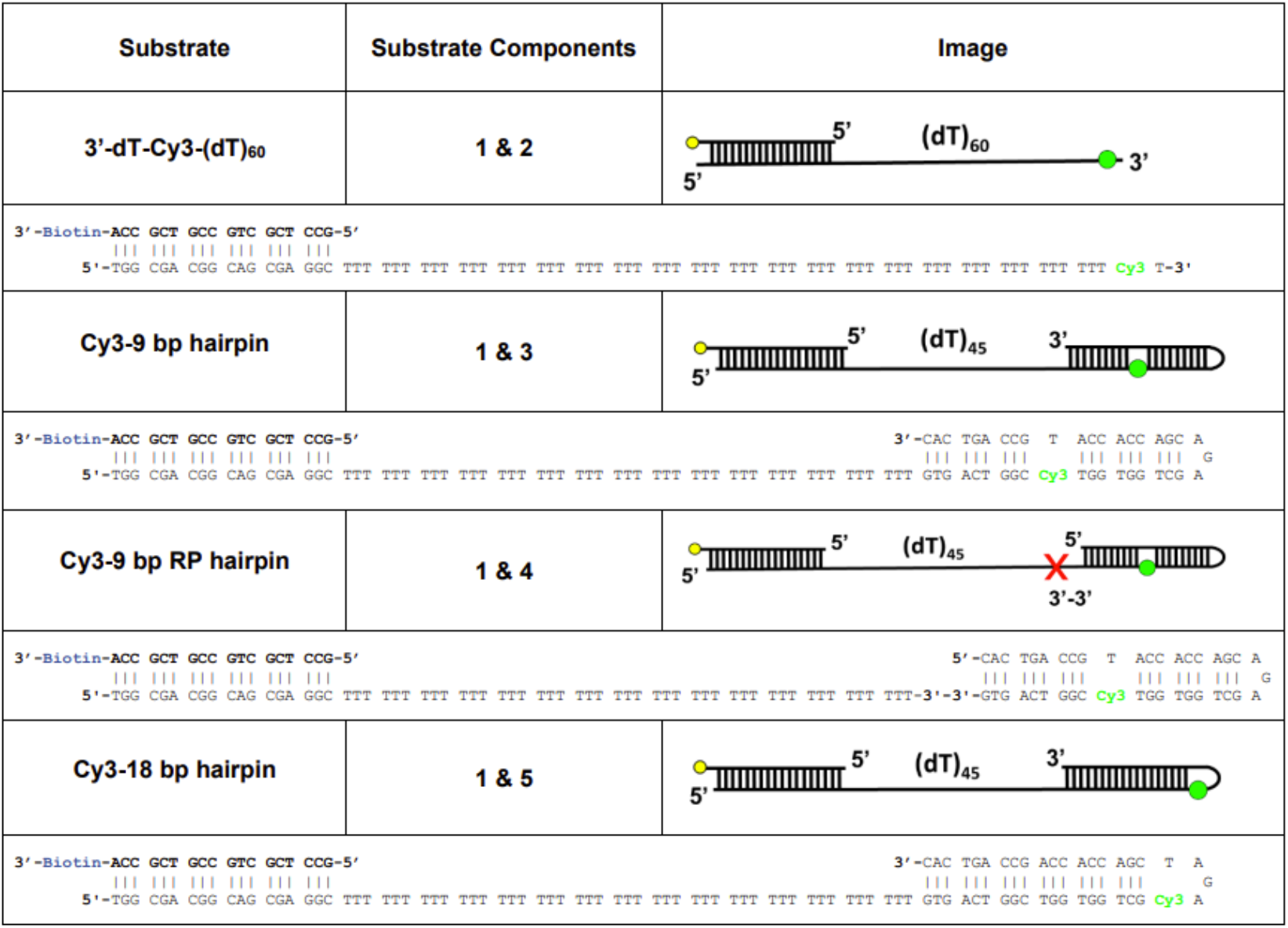
Single Molecule TIRF Substrates and Base Pairing

**Fig. S1.**
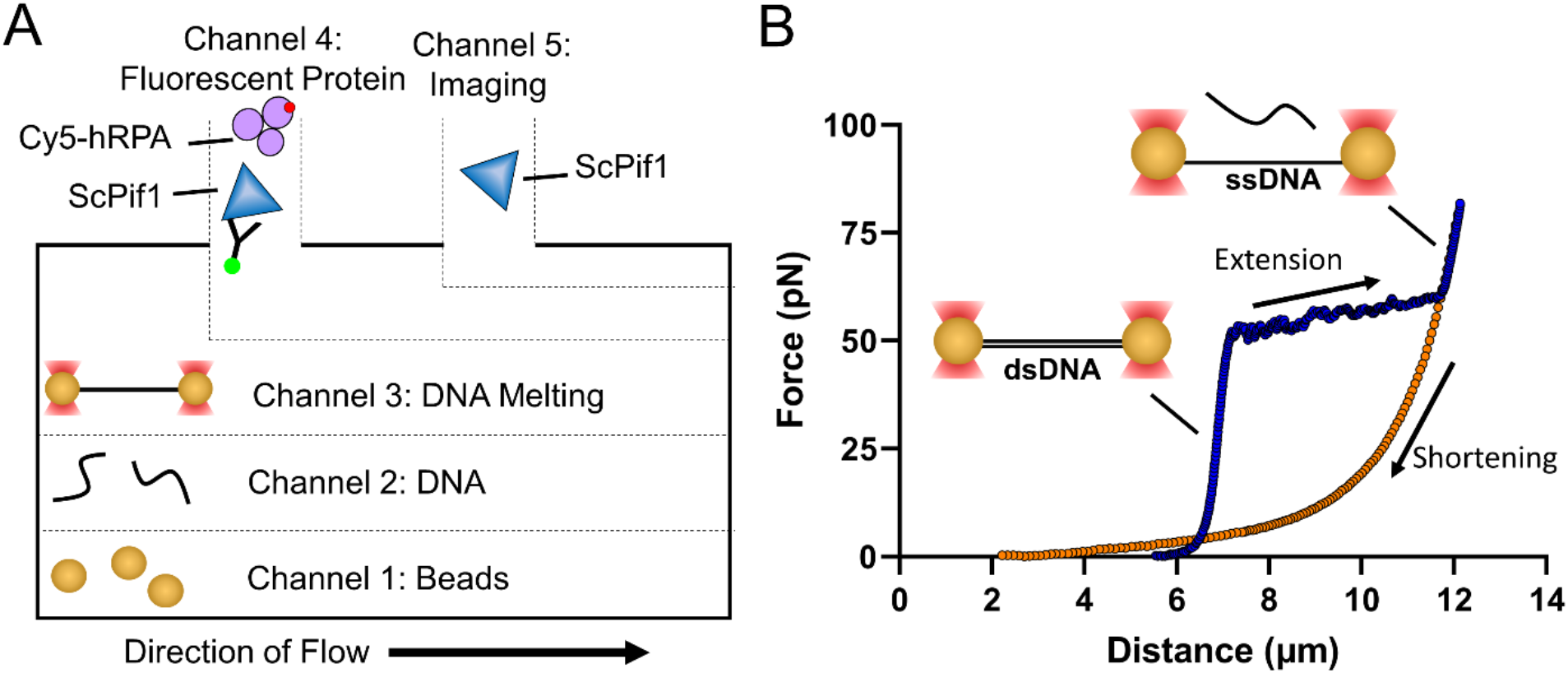
Flow chamber and single-stranded DNA formation. **A)** Schematic of the Lumicks flow cell used in the optical tweezers experiments. Channel 1 contains polystyrene beads (4.89 μm diameter) coated with streptavidin, while Channel 2 contains biotinylated dsDNA (20,452 bp). Channel 3 is where force was applied to dsDNA to form ssDNA. Channels 4 and 5 can have Cy5-hRPA, Pif1, or Pif1-N-His + DyL550-IgG, and imaging was performed exclusively in Channel 5. **B)** Force-extension curves were performed to confirm the formation of ssDNA after the dsDNA tether was held at a constant distance of 12.3 μm for 2 seconds.

**Fig. S2.**
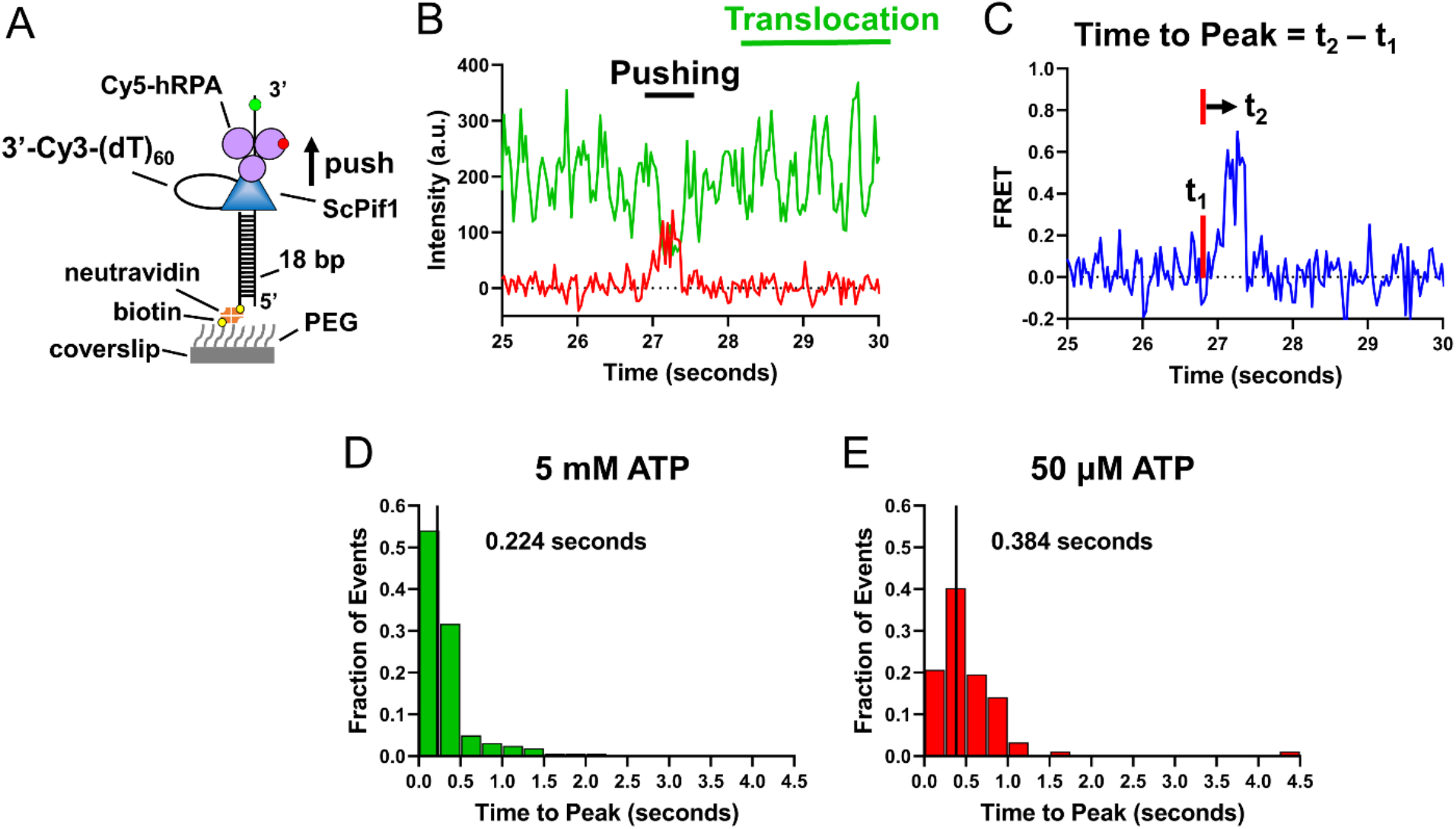
Time-to-Peak assay reveals hRPA pushing by Pif1 is [ATP] dependent. **A)** Pif1 pushing Cy5-hRPA off a Cy3-(dT)_60_ ssDNA substrate with a 5’ biotinylated 18 bp dsDNA handle immobilized on the surface of a neutravidin-treated biotinylated-PEG coverslip. **B)** Time trajectory from a smTIRF experiment for Pif1 pushing a Cy5-hRPA off a 3’ Cy3-labeled (dT)_60_ DNA. The Cy3 time trajectory is green, and the Cy5 time trajectory is red. The anti-correlated behavior indicates the occurrence of a FRET event. **C)** Calculated FRET signal from the emission intensities presented in B. The time-to-peak value is defined by the time elapsed while an asymmetric spike rises from the baseline FRET efficiency of the trajectory to the maximum FRET of the spike. **D and E)** Time-to-peak histograms for Cy5-hRPA being pushed along Cy3-(dT)_60_ ssDNA by Pif1 at **D)** 5 mM ATP (median = 0.224 (n = 161 events)) and **E)** 50 μM ATP (median = 0.384 (n = 92 events)).

**Fig. S3.**
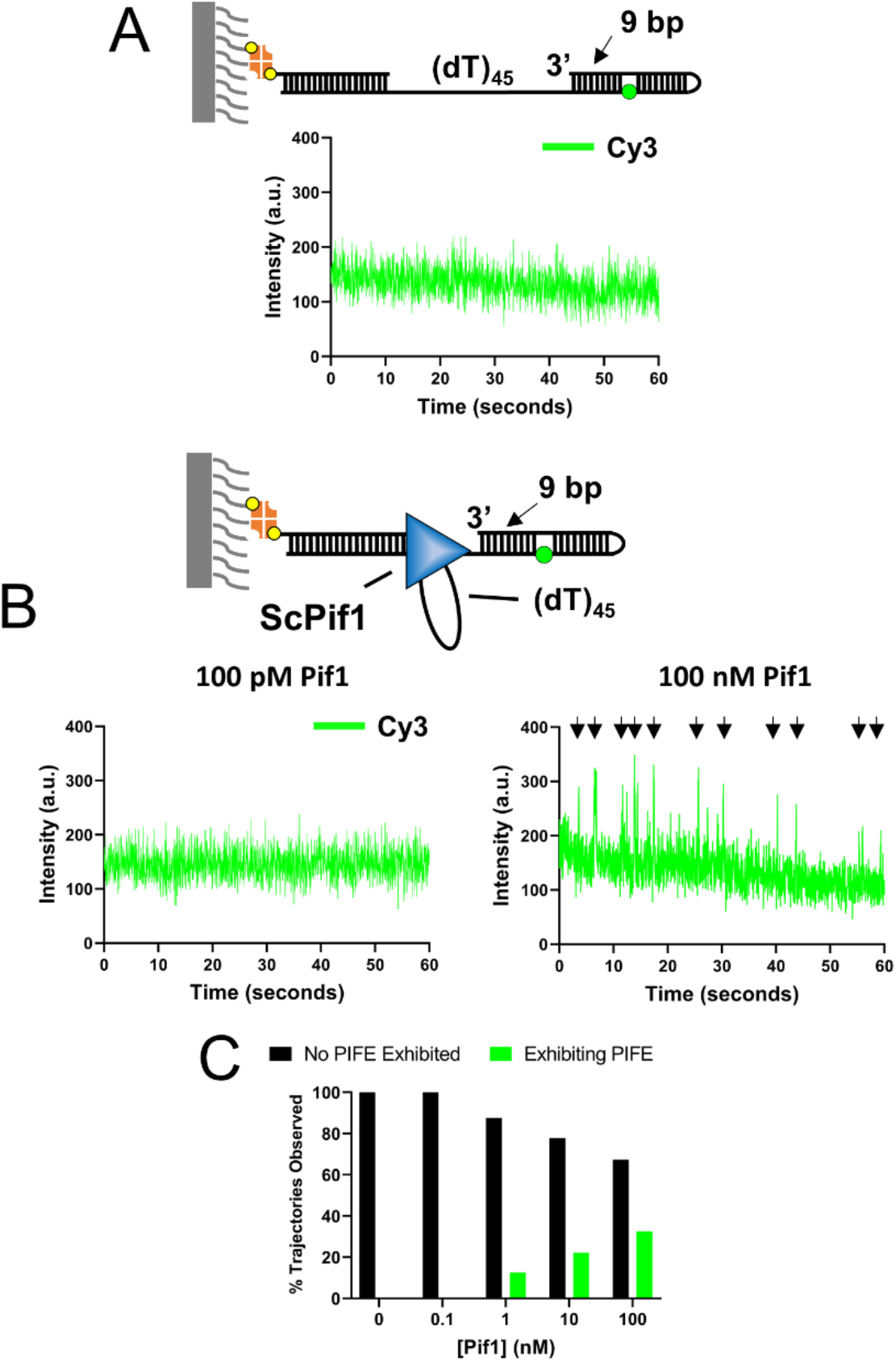
Disruption of a Cy3-9 bp hairpin is Pif1 concentration dependent. **A)** smTIRF DNA substrate with a (dT)_45_ ssDNA region at the 5’ end of an 18 bp hairpin with a Cy3 fluorophore present in the backbone 9 bp from the ss/dsDNA bound to a smTIRF slide. smTIRFM trajectory depicts the Cy3 emission of the Cy3-9 bp hairpin construct with no added Pif1. No enhancements in Cy3 fluorescence were observed (0 mM Pif1 (n = 0, N = 820). **B)** (Left) On the Cy3-9 bp hairpin construct from A, no enhancements in Cy3 occur in the presence of 100 pM Pif1 and 5 mM ATP (n = 0, N = 180). (Right) Repetitive PIFE events occur in the Cy3 emission signal when the experiments are performed at [Pif1] ≥ 1 nM. Black arrows indicated PIFE events. **C)** Percentage of smTIRF trajectories recorded on the Cy3-9 bp hairpin construct that exhibit PIFE events at [ATP] = 5 mM and increasing [Pif1] (100 pM Pif1 (n = 0, N = 180), 1 nM (n = 22, N = 176), 10 nM (n = 30, N = 135), 100 nM (n = 58, N = 178)). Data for DNA only (0 nM Pif1, (n = 0, N = 820)) are shown as reference.

## References

1. R. Chen, M. S. Wold, Replication protein A: single-stranded DNA’s first responder: dynamic DNA-interactions allow replication protein A to direct single-strand DNA intermediates into different pathways for synthesis or repair. Bioessays 36, 1156–1161 (2014).

2. M. S. Wold, Replication protein A: a heterotrimeric, single-stranded DNA-binding protein required for eukaryotic DNA metabolism. Annu Rev Biochem 66, 61–92 (1997).

3. B. Nguyen et al., Diffusion of human replication protein A along single-stranded DNA. J Mol Biol 426, 3246–3261 (2014).

4. N. Pokhrel et al., Dynamics and selective remodeling of the DNA-binding domains of RPA. Nat Struct Mol Biol 26, 129–136 (2019).

5. F. E. Kemmerich et al., Force regulated dynamics of RPA on a DNA fork. Nucleic Acids Res 44, 5837–5848 (2016).

6. R. Roy, A. G. Kozlov, T. M. Lohman, T. Ha, SSB protein diffusion on single-stranded DNA stimulates RecA filament formation. Nature 461, 1092–1097 (2009).

7. R. Zhou et al., SSB functions as a sliding platform that migrates on DNA via reptation. Cell 146, 222–232 (2011).

8. K. S. Lee et al., Ultrafast redistribution of E. coli SSB along long single-stranded DNA via intersegment transfer. J Mol Biol 426, 2413–2421 (2014).

9. A. G. Kozlov, T. M. Lohman, Kinetic mechanism of direct transfer of Escherichia coli SSB tetramers between single-stranded DNA molecules. Biochemistry 41, 11611–11627 (2002).

10. S. Kunzelmann, C. Morris, A. P. Chavda, J. F. Eccleston, M. R. Webb, Mechanism of interaction between single-stranded DNA binding protein and DNA. Biochemistry 49, 843–852 (2010).

11. B. Gibb et al., Concentration-dependent exchange of replication protein A on single-stranded DNA revealed by single-molecule imaging. PLoS One 9, e87922 (2014).

12. J. E. Sokoloski, A. G. Kozlov, R. Galletto, T. M. Lohman, Chemo-mechanical pushing of proteins along single-stranded DNA. Proc Natl Acad Sci U S A 113, 6194–6199 (2016).

13. S. Hormeno et al., Human HELB is a processive motor protein that catalyzes RPA clearance from single-stranded DNA. Proc Natl Acad Sci U S A 119, e2112376119 (2022).

14. E. Antony, T. M. Lohman, Dynamics of E. coli single stranded DNA binding (SSB) protein-DNA complexes. Semin Cell Dev Biol 86, 102–111 (2019).

15. R. Zhou, J. Zhang, M. L. Bochman, V. A. Zakian, T. Ha, Periodic DNA patrolling underlies diverse functions of Pif1 on R-loops and G-rich DNA. Elife 3, e02190 (2014).

16. B. Nguyen, M. A. Ciuba, A. G. Kozlov, M. Levitus, T. M. Lohman, Protein Environment and DNA Orientation Affect Protein-Induced Cy3 Fluorescence Enhancement. Biophys J 117, 66–73 (2019).

17. H. Hwang, H. Kim, S. Myong, Protein induced fluorescence enhancement as a single molecule assay with short distance sensitivity. Proc Natl Acad Sci U S A 108, 7414–7418 (2011).

18. R. Galletto, E. J. Tomko, Translocation of Saccharomyces cerevisiae Pif1 helicase monomers on single-stranded DNA. Nucleic Acids Res 41, 4613–4627 (2013).

19. S. P. Singh, A. Soranno, M. A. Sparks, R. Galletto, Branched unwinding mechanism of the Pif1 family of DNA helicases. Proc Natl Acad Sci U S A 116, 24533–24541 (2019).

20. S. P. Singh, K. N. Koc, J. L. Stodola, R. Galletto, A Monomer of Pif1 Unwinds Double-Stranded DNA and It Is Regulated by the Nature of the Non-Translocating Strand at the 3’-End. J Mol Biol 428, 1053–1067 (2016).

21. M. N. Dessinges et al., Stretching single stranded DNA, a model polyelectrolyte. Phys Rev Lett 89, 248102 (2002).

22. A. Bosco, J. Camunas-Soler, F. Ritort, Elastic properties and secondary structure formation of single-stranded DNA at monovalent and divalent salt conditions. Nucleic Acids Res 42, 2064–2074 (2014).

23. M. A. Sparks, S. P. Singh, P. M. Burgers, R. Galletto, Complementary roles of Pif1 helicase and single stranded DNA binding proteins in stimulating DNA replication through G-quadruplexes. Nucleic Acids Res 47, 8595–8605 (2019).

24. W. C. Griffin, J. Gao, A. K. Byrd, S. Chib, K. D. Raney, A biochemical and biophysical model of G-quadruplex DNA recognition by positive coactivator of transcription 4. J Biol Chem 292, 9567–9582 (2017).

25. D. Dahan et al., Pif1 is essential for efficient replisome progression through lagging strand G-quadruplex DNA secondary structures. Nucleic Acids Res 46, 11847–11857 (2018).

26. M. A. Sparks, P. M. Burgers, R. Galletto, Pif1, RPA, and FEN1 modulate the ability of DNA polymerase delta to overcome protein barriers during DNA synthesis. J Biol Chem 295, 15883–15891 (2020).

27. M. E. Douglas, J. F. X. Diffley, Budding yeast Rap1, but not telomeric DNA, is inhibitory for multiple stages of DNA replication in vitro. Nucleic Acids Res 49, 5671–5683 (2021).

28. G. D. Schauer et al., Replisome bypass of a protein-based R-loop block by Pif1. Proc Natl Acad Sci U S A 117, 30354–30361 (2020).

29. P. Baumann, S. C. West, The human Rad51 protein: polarity of strand transfer and stimulation by hRP-A. EMBO J 16, 5198–5206 (1997).

30. S. Awate, R. M. Brosh, Jr., Interactive Roles of DNA Helicases and Translocases with the Single-Stranded DNA Binding Protein RPA in Nucleic Acid Metabolism. Int J Mol Sci 18 (2017).

31. M. M. Cox, Motoring along with the bacterial RecA protein. Nat Rev Mol Cell Biol 8, 127–138 (2007).

32. S. L. Lusetti, M. M. Cox, The bacterial RecA protein and the recombinational DNA repair of stalled replication forks. Annu Rev Biochem 71, 71–100 (2002).

33. S. C. Kowalczykowski, D. A. Dixon, A. K. Eggleston, S. D. Lauder, W. M. Rehrauer, Biochemistry of homologous recombination in Escherichia coli. Microbiol Rev 58, 401–465 (1994).

34. R. M. Brosh, Jr. et al., Functional and physical interaction between WRN helicase and human replication protein A. J Biol Chem 274, 18341–18350 (1999).

35. D. Bagchi et al., Single molecule kinetics uncover roles for E. coli RecQ DNA helicase domains and interaction with SSB. Nucleic Acids Res 46, 8500–8515 (2018).

36. T. M. Lohman, E. J. Tomko, C. G. Wu, Non-hexameric DNA helicases and translocases: mechanisms and regulation. Nat Rev Mol Cell Biol 9, 391–401 (2008).

37. K. M. Brendza et al., Autoinhibition of Escherichia coli Rep monomer helicase activity by its 2B subdomain. Proc Natl Acad Sci U S A 102, 10076–10081 (2005).

38. W. Cheng, J. Hsieh, K. M. Brendza, T. M. Lohman, E. coli Rep oligomers are required to initiate DNA unwinding in vitro. J Mol Biol 310, 327–350 (2001).

39. K. S. Lee, H. Balci, H. Jia, T. M. Lohman, T. Ha, Direct imaging of single UvrD helicase dynamics on long single-stranded DNA. Nat Commun 4, 1878 (2013).

40. A. Niedziela-Majka, M. A. Chesnik, E. J. Tomko, T. M. Lohman, Bacillus stearothermophilus PcrA monomer is a single-stranded DNA translocase but not a processive helicase in vitro. J Biol Chem 282, 27076–27085 (2007).

41. N. K. Maluf, J. A. Ali, T. M. Lohman, Kinetic mechanism for formation of the active, dimeric UvrD helicase-DNA complex. J Biol Chem 278, 31930–31940 (2003).

42. N. K. Maluf, C. J. Fischer, T. M. Lohman, A Dimer of Escherichia coli UvrD is the active form of the helicase in vitro. J Mol Biol 325, 913–935 (2003).

43. C. J. Fischer, N. K. Maluf, T. M. Lohman, Mechanism of ATP-dependent translocation of E.coli UvrD monomers along single-stranded DNA. J Mol Biol 344, 1287–1309 (2004).

44. B. Nguyen, Y. Ordabayev, J. E. Sokoloski, E. Weiland, T. M. Lohman, Large domain movements upon UvrD dimerization and helicase activation. Proc Natl Acad Sci U S A 114, 12178–12183 (2017).

45. A. Chadda et al., Mycobacterium tuberculosis DNA repair helicase UvrD1 is activated by redox-dependent dimerization via a 2B domain cysteine. Proc Natl Acad Sci U S A 119 (2022).

46. B. Nguyen, M. K. Shinn, E. Weiland, T. M. Lohman, Regulation of E. coli Rep helicase activity by PriC. J Mol Biol 433, 167072 (2021).

47. Y. A. Ordabayev, B. Nguyen, A. G. Kozlov, H. Jia, T. M. Lohman, UvrD helicase activation by MutL involves rotation of its 2B subdomain. Proc Natl Acad Sci U S A 116, 16320–16325 (2019).

48. Y. A. Ordabayev, B. Nguyen, A. Niedziela-Majka, T. M. Lohman, Regulation of UvrD Helicase Activity by MutL. J Mol Biol 430, 4260–4274 (2018).

49. L. T. Chisty et al., Monomeric PcrA helicase processively unwinds plasmid lengths of DNA in the presence of the initiator protein RepD. Nucleic Acids Res 41, 5010–5023 (2013).

50. S. Barranco-Medina, R. Galletto, DNA binding induces dimerization of Saccharomyces cerevisiae Pif1. Biochemistry 49, 8445–8454 (2010).

